# A Hierarchy-aware Gene Exploration Platform for Multi-layered Toxicogenomic Analysis: A Case Study on Acetaminophen-induced Hepatotoxicity

**DOI:** 10.64898/2026.04.10.717684

**Authors:** MinJung Kim, YaXuan Cui, Myeong Gyu Kim

## Abstract

**Background:** The interpretation of high-dimensional transcriptomic data remains a major challenge in mechanistic toxicology and drug safety assessment. Conventional clustering approaches based solely on expression profiles often fail to capture intrinsic biological relationships among genes, limiting interpretability and downstream analysis.

**Methods:** We developed a hierarchy-aware gene exploration platform that integrates structured biological knowledge from the HUGO Gene Nomenclature Committee (HGNC). The core of the framework is a similarity kernel based on a single-step hyperdiffusion formulation (*HKH*^⊤^), which embeds gene family hierarchy into the similarity space. The platform is implemented as an interactive web application supporting Uniform Manifold Approximation and Projection (UMAP) visualization, Leiden clustering, functional enrichment analysis, and hierarchy-based gene recommendation.

**Results:** Applied to a transcriptomic dataset of acetaminophen-induced acute liver failure (APAP–ALF), the proposed approach achieved a 33.8-fold improvement in functional coherence compared to an expression-only baseline. The hierarchy-aware embedding produced compact and biologically consistent clusters, enabling identification of key toxicological modules, including disruption of RNA processing, extracellular matrix remodeling, and impairment of lipid transport. In addition, the framework detected small but highly significant regulatory modules associated with epigenetic reprogramming.

**Conclusion:** By incorporating biological hierarchy into gene similarity, the proposed platform enhances the interpretability of transcriptomic analysis and enables structured exploration of functional relationships. This approach provides a practical framework for mechanistic insight generation and supports more transparent and reproducible analysis in toxicogenomics.

**Availability:** The web application is freely available at https://hgncgeneexplorer.streamlit.app/.

## 1 Introduction

The rapid advancement of high-throughput transcriptomics has enabled the generation of vast amounts of gene expression data.[1] However, a significant bottleneck remains in the functional interpretation of these data.[2] Traditional analytical pipelines often rely on clustering methods based solely on expression profiles, which frequently fail to capture the underlying biological relationships and evolutionary lineages of human genes. Furthermore, most existing tools provide static outputs that lack the interactive depth required for real-time hypothesis generation and regulatory decision-making.

To address these limitations, we present a hierarchy-aware gene exploration platform that integrates structured biological knowledge from the HUGO Gene Nomenclature Committee (HGNC).[3] The core of our approach is a biologically informed similarity kernel (*HKH*^⊤^), which embeds gene family hierarchy directly into the similarity space. By redefining gene–gene relationships through hierarchical lineage, our framework enables the identification of functionally coherent modules that are consistent with established gene families, rather than relying solely on expression proximity.

The platform is implemented as a publicly accessible web application (https://hgncgeneexplorer.streamlit.app/), supporting interactive analysis through UMAP visualization, Leiden clustering, functional enrichment, and hierarchy-based gene exploration. We evaluate the proposed framework using a transcriptomic dataset of acetaminophen (APAP)-induced acute liver failure (APAP-ALF), demonstrating its ability to recover biologically meaningful structures from complex gene expression data.

By bridging raw transcriptomic signals with structured biological nomenclature, this work provides a practical and interpretable framework for toxicogenomic analysis and mechanistic insight generation.

## 2 Methods

### 2.1 Data Sources and Preprocessing

The proposed analytical pipeline requires three primary data inputs: (i) HGNC gene metadata (`hgnc_complete_set.txt`), (ii) HGNC gene–family hierarchy closure (`hierarchy_closure.csv`), containing parent–child family relationships with shortest-path distance *d*, and (iii) a curated experimental gene list with associated numeric values (e.g., fold change). The overall computational workflow of the proposed framework is illustrated in Figure 1. The workflow consists of data integration, ontology mapping, hierarchy-aware similarity construction, downstream clustering and visualization, and a recommendation module.

**Figure 1.**
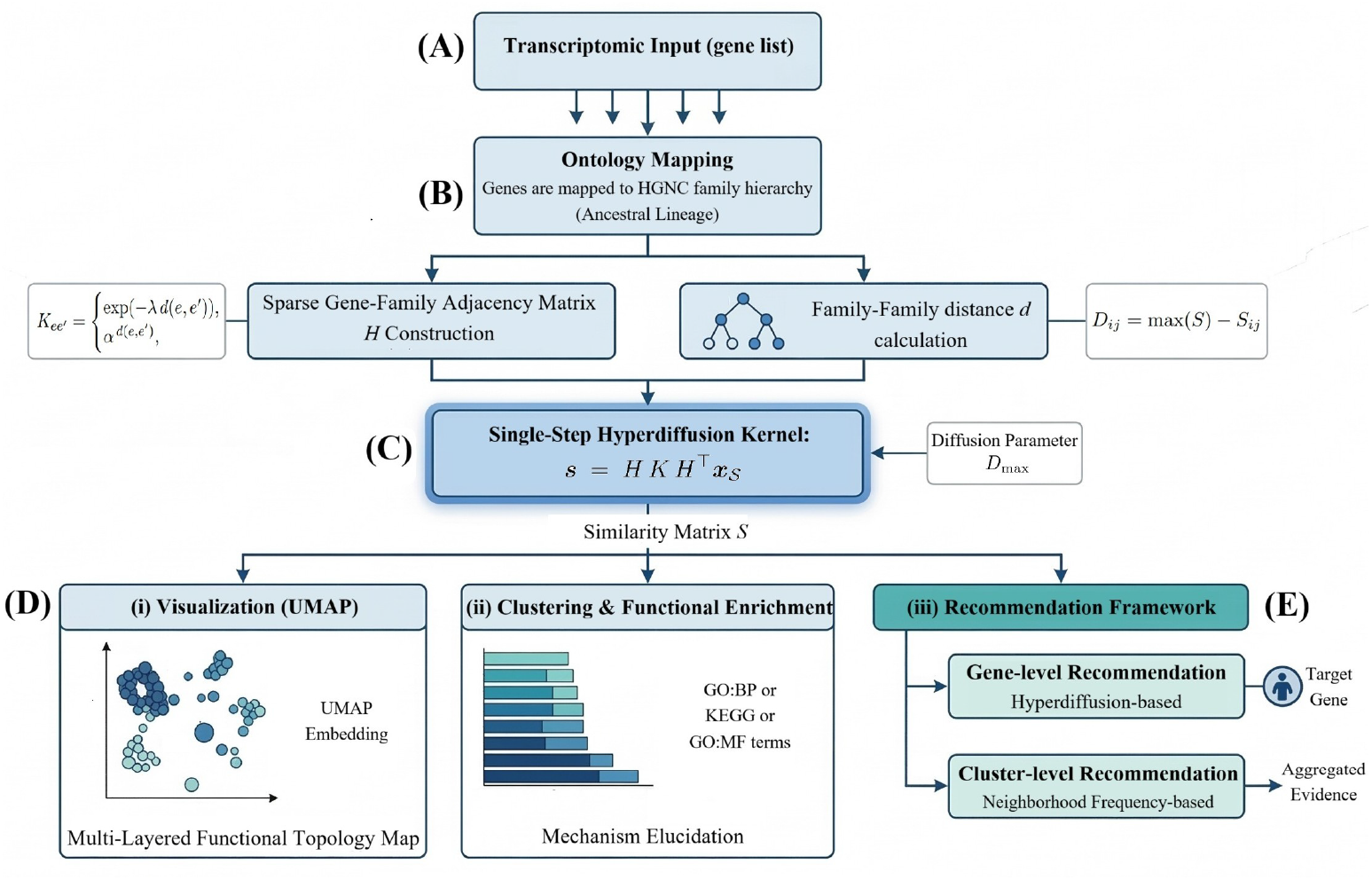
Overview of the HGNC hierarchy-aware computational framework. (A) Data integration, (B) HGNC ontology mapping, (C) hierarchy-aware similarity construction via hyperdiffusion, (D) downstream analysis including UMAP visualization and clustering, and (E) recommendation framework.

#### HGNC resources

HGNC data were obtained from `genenames.org` (HUGO Gene Nomenclature Committee),[3] including (i) `hgnc_complete_set`, containing approved human gene symbols and metadata, and (ii) `hierarchy_closure.csv`, providing transitive family relationships with shortest-path distances. We used the HGNC quarterly snapshot (2025-03-18) to ensure reproducibility.

#### Experimental Dataset

To demonstrate the regulatory utility of the platform, we utilized a transcriptomic dataset derived from APAP-ALF (GEO Series: GSE74000).[4] This dataset contains liver tissue gene expression profiles from APAP-ALF patients and healthy human controls. For our analysis, the fold change (ALF vs. Healthy) was mapped to the tool’s numeric value attribute (labeled as Count).

#### Preprocessing

Gene symbols were standardized to upper case and mapped to HGNC identifiers using exact matching, with fallback matching via `alias_symbol` and `prev_ symbol`. Unmapped genes were excluded.

Genes without family membership were excluded by default, although optional inclusion is supported. If a numeric attribute is provided, values may be transformed as 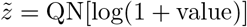 for optional downstream weighting.

### 2.2 Hierarchical Prior and Gene Similarity

#### Hierarchy-aware kernel

To encode biological lineage relationships, we construct a family–family proximity kernel *K ∈ ℝ*^*m×m*^ using hierarchy closure distances. For pairs with distance *d ≤ D*_max_:

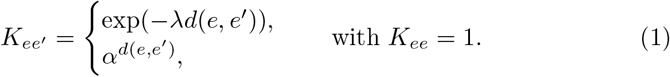

#### Gene similarity matrix

Let *A ∈ ℝ*^*n×m*^ denote the gene–family incidence matrix. To mitigate bias from large families, we apply normalization *A*_*ie*_ = 1*/*|*e*|.

The gene–gene similarity matrix is defined as:

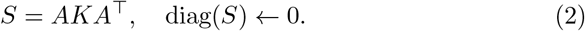

Optionally, count-based weighting can be applied:

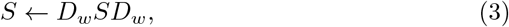

where *D*_*w*_ = diag(**w**).

### 2.3 Graph Construction and Clustering

For computational efficiency and to emphasize local structure, the similarity matrix *S* is sparsified using row-wise top-*k* filtering, retaining the strongest connections per gene. The matrix is then symmetrized to form an undirected graph.

Clustering is performed using the Leiden algorithm,[5] with edge weights derived from *S*. We employ the `RBConfigurationVertexPartition` objective, allowing resolution-controlled clustering to identify functionally coherent modules.

### 2.4 Two-dimensional Embedding

To visualize the hierarchy-aware genomic space, the similarity matrix is converted into a distance matrix:

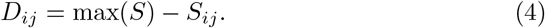

Pairs without shared lineage within the hierarchy radius are assigned the maximum distance. Low-dimensional embedding is performed using UMAP with a `precomputed` metric.[6] Hyperparameters such as *n_neighbors* and *min_dist* are selected relative to dataset size.

### 2.5 Functional Enrichment Analysis

To characterize biological relevance of identified clusters, we perform enrichment analysis using the `g:Profiler` API. [7] For each cluster, gene symbols are queried against the human reference database.

Statistical overrepresentation (*p <* 0.05) is evaluated across: (i) Gene Ontology Biological Process (GO:BP), (ii) Gene Ontology Molecular Function (GO:MF),[8] and (iii) KEGG pathways.[9]

### 2.6 Recommendation Framework

We propose a hierarchy-aware recommendation framework that prioritizes candidate genes based on HGNC family relationships. The framework consists of two complementary strategies: (i) gene-level recommendation, which propagates signals from one or more seed genes, and (ii) cluster-level recommendation, which aggregates evidence across a group of curated genes.The overall mechanism of hierarchy-based propagation is illustrated in Figure 2.

**Figure 2.**
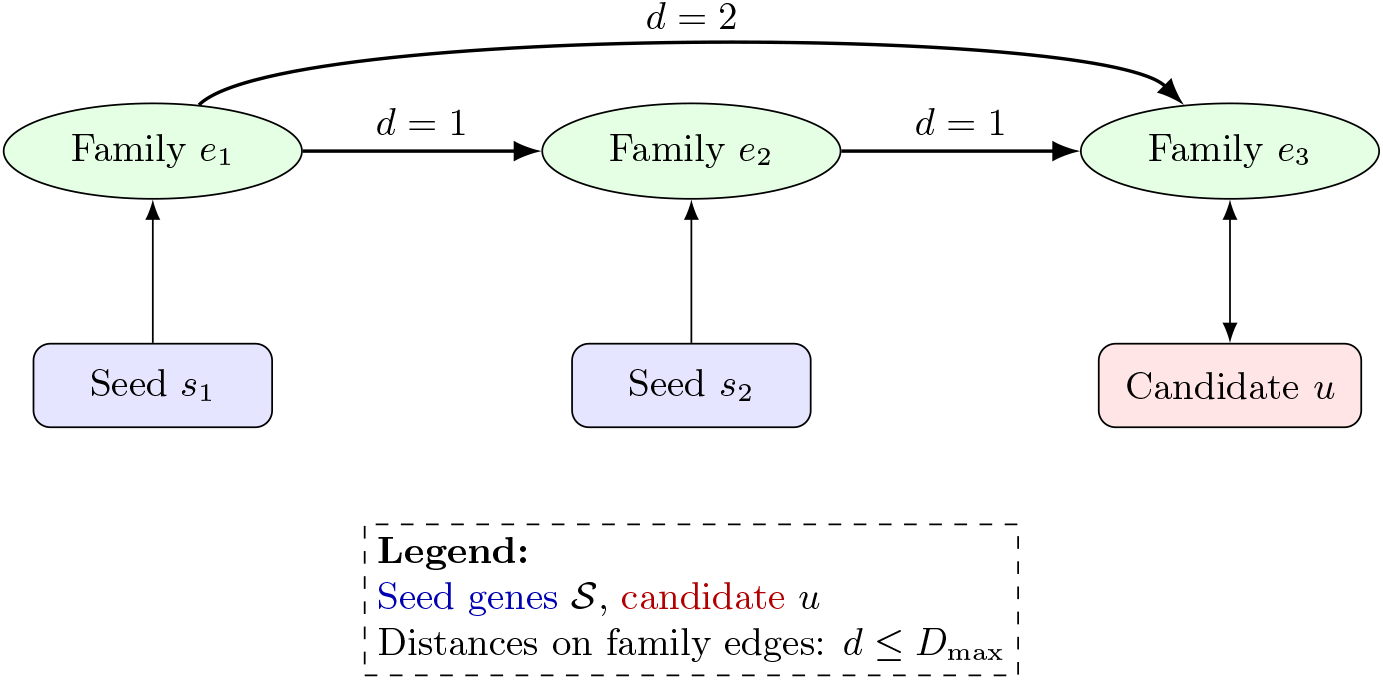
Family-hierarchy–aware recommendation. Seeds (*s*_1_, *s*_2_) diffuse through family proximity to score candidate *u*.

Both approaches share a common propagation backbone:

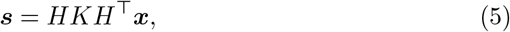

where *H∈ ℝ*^*n×m*^ is the gene–family incidence matrix, *K∈ ℝ*^*m×m*^ is the hierarchy-aware kernel, and ***x*** is the seed indicator vector. This formulation enables memory-efficient computation without explicitly constructing a dense *n × n* similarity matrix.

#### 2.6.1 Gene-level Recommendation

Given a seed set *𝒮* ⊆ {1, …, *n*}, we define an indicator vector *x*_*𝒮*_ ∈ {0, 1}^*n*^ such that (***x***_*S*_)_*i*_ = 1 if *i ∈ 𝒮* and 0 otherwise. The recommendation score is obtained via:

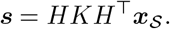

This operation can be interpreted as a single-step diffusion process across the gene–family hierarchy. Scores are min–max normalized to [0, 1], and genes with zero scores (i.e., no hierarchical connection within *D*_max_) are discarded. Candidate genes are selected from the HGNC universe excluding the seed set and curated input list.

##### Algorithm 1

Gene-level Recommendation (memory-efficient)

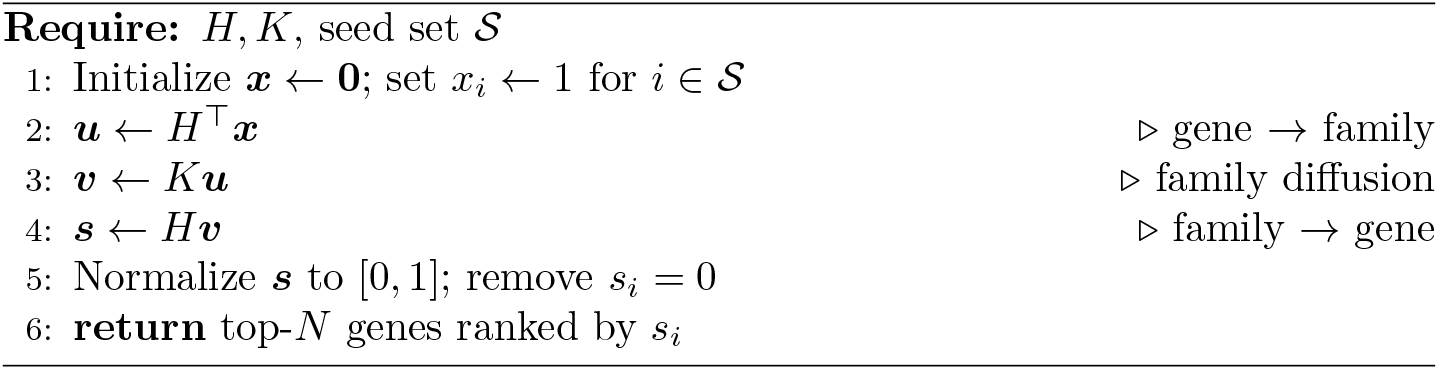

#### 2.6.2 Cluster-level Recommendation

For a selected cluster *c* with curated gene set *𝒮*_*c*_, we compute the base propagation score 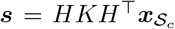 . To incorporate both structural proximity and collective evidence, we define the following quantities for each candidate gene *u*:

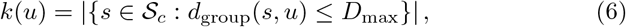

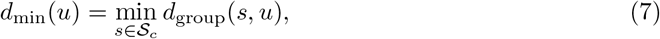

where *d*_group_ denotes the minimum family-level distance in the HGNC hierarchy.

The final recommendation score is defined as:

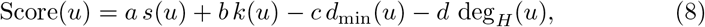

where *a, b, c, d* ≥0 are user-adjustable weights and deg_*H*_ (*u*) is the number of families gene *u* belongs to.

The linear formulation enables interpretable integration of proximity, frequency, and specificity. The hub penalty term deg_*H*_ (*u*) reduces bias toward highly connected genes that may otherwise dominate recommendations. Candidates with *s*(*u*) = 0 are excluded, and remaining genes are ranked in descending order.

### 2.7 Platform Architecture and Implementation

The system is implemented using a modular architecture:

- **Data engine:** handles HGNC and experimental datasets using `Pandas` and `NumPy` .
- **Computational core:** performs similarity computation and clustering using sparse matrices and `Leidenalg` .
- **Interface layer:** built with `Streamlit` for interactive user control.
- **Visualization:** implemented using `Plotly` and `UMAP-learn` .

### 2.8 Evaluation Protocol

To assess functional coherence, we compare the hierarchy-aware model with an expression-only baseline.

For each cluster, enrichment analysis is performed, and coherence is quantified as:

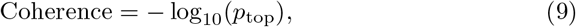

where *p*_top_ is the most significant enrichment result.

Only clusters with *n ≥* 4 genes are included to ensure robustness.

### 2.9 Parameter Settings and Oversmoothing

Default parameters are set as follows: *D*_max_ = 3; for exponential decay, *λ ∈* [0.7, 1.0]; for power decay, *α ∈* [0.5, 0.8].

Larger values of *D*_max_ or weaker decay can lead to *oversmoothing*, where excessive propagation reduces discriminative structure and causes genes to become indistinguishable in the embedding space. These parameters should therefore be tuned to balance local specificity and global connectivity.

## 3 Results

### 3.1 Workflow of the HGNC Hierarchy-aware Gene Exploration System

Figure 3 illustrates the interactive workflow of the proposed platform. Users can upload curated gene lists and configure parameters through an intuitive sidebar interface (Figure 3(a)). Gene queries can be performed using symbols or HGNC IDs (Figure 3(b)), with results visualized directly on the UMAP embedding (Figure 3(c)).

**Figure 3.**
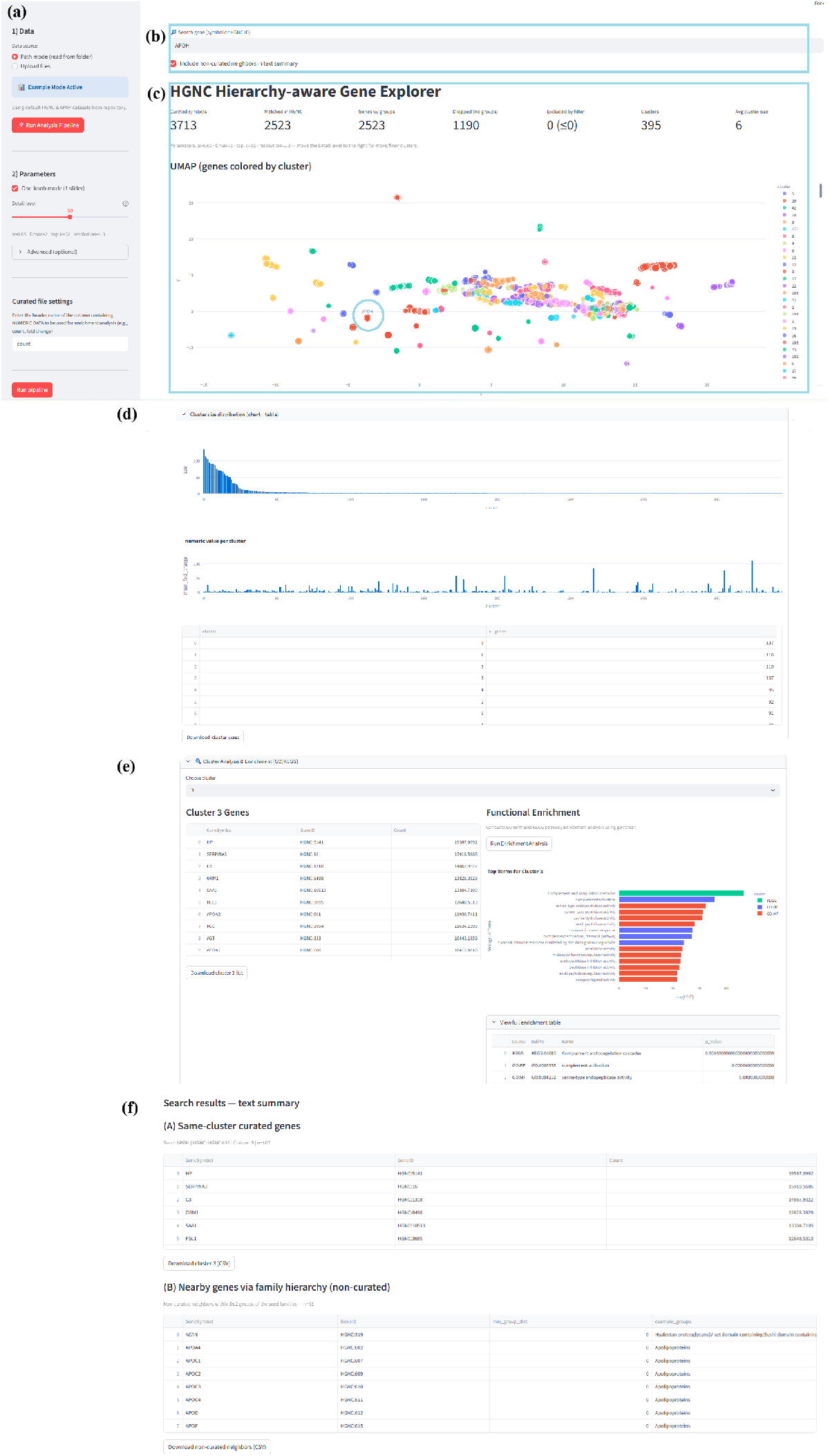
Overview of the HGNC hierarchy-aware gene exploration workflow.

The system provides cluster-level summaries, including size distributions and numerical statistics, which can be interactively explored and downloaded (Figure 3(d)). Users can further investigate individual clusters through gene lists and functional enrichment analysis (Figure 3(e)).

In addition, an integrated search module provides contextual interpretation of queried genes by displaying same-cluster curated genes and hierarchy-based neighboring genes, enabling exploration across both local cluster membership and broader biological relationships (Figure 3(f)).

### 3.2 Comparison of Baseline and Hierarchy-aware Embeddings

To evaluate the structural impact of incorporating the HGNC hierarchy, we compared the proposed model with a structure-agnostic baseline. As shown in Figure 4(a), the baseline UMAP exhibits a broadly distributed embedding in which points are densely intermingled across the space, with no clearly defined boundaries or localized group structure.

**Figure 4.**
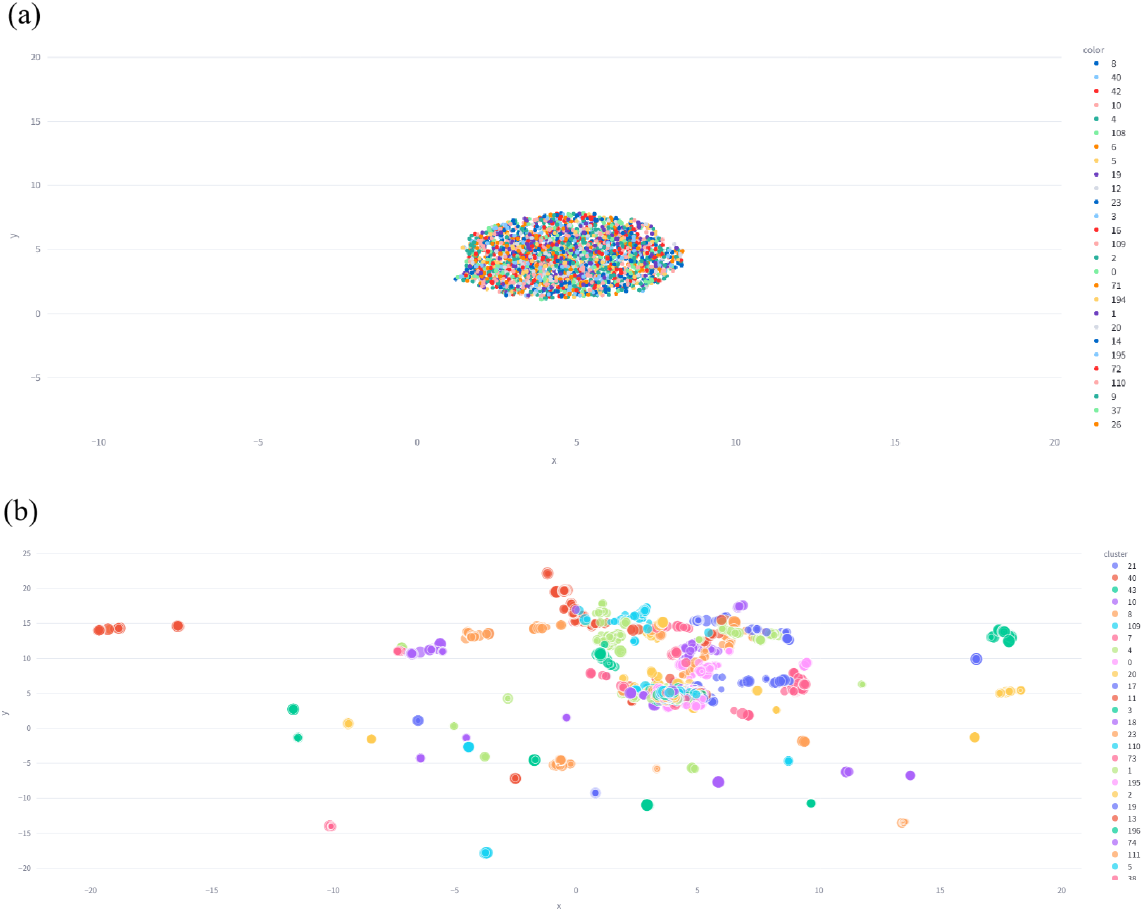
Comparison of baseline and hierarchy-aware embeddings of genes. UMAP visualization of genes from the APAP–ALF dataset colored by cluster labels. (a) Baseline UMAP constructed from structure-agnostic features. (b) Hierarchy-aware UMAP constructed using the HGNC gene family hierarchy.

In contrast, the hierarchy-aware embedding (Figure 4(b)) shows more coherent and localized groupings, with reduced overlap and more distinct separation between regions. Genes within these regions tend to share functional annotations, suggesting improved organization of biologically related genes in the embedding space. These observations indicate that incorporating the HGNC hierarchy leads to a more structured representation compared to the baseline.

### 3.3 Data Integration and Hierarchy-aware Clustering

We applied the proposed framework to a transcriptomic dataset of APAP-ALF(GEO: GSE74000). [4] Among 3,713 curated gene symbols, 2,523 genes were successfully mapped to HGNC and assigned to functional clusters. The resulting hierarchy-aware similarity structure partitioned the gene space into 396 clusters.

Unlike conventional expression-based clustering, the proposed method preserves biological lineage relationships defined by the HGNC hierarchy. As a result, genes belonging to the same functional family remain grouped even when their expression magnitudes differ substantially.

### 3.4 Quantitative Evaluation of Functional Coherence

We next evaluated the biological relevance of the identified clusters using an ablation study against an expression-only baseline (Table 1). The proposed model achieved an average functional coherence score of 17.04, representing a 33.8-fold increase over the baseline score of 0.50.

**Table 1.** Ablation Analysis: Impact of HGNC Hierarchy on Cluster Coherence.

| Configuration | Hierarchy ($H$) | Avg. Coherence<sup>a</sup> | Improvement |
| --- | --- | --- | --- |
| Baseline (Expression-only) | No | 0.50 | 1.0× |
| **Proposed** | **Yes** | **17.04** | **33.8×** |

<sup>a</sup> Average functional coherence: $\text{mean} - \log_{10}(P\text{-value})$ for clusters with $n \geq 4$ .

This improvement corresponds to a substantial gain in statistical significance, with the proposed model identifying highly enriched biological processes (*P≈* 10^*−*17^) compared to the baseline (*P ≈* 0.31). These results indicate that incorporating the HGNC hierarchy significantly enhances the recovery of biologically meaningful gene modules.

^a^ Average functional coherence: mean *−* log_10_(*P* -value) for clusters with *n ≥* 4.

### 3.5 Identification of Toxicological Mechanisms

The platform identified three primary functional modules characterizing the hepatic response to APAP insult (Figure 5):

- **Intracellular RNA Processing (Cluster 6)**: This module (*n* = 94) showed significant enrichment in the spliceosome pathway (KEGG:03040, *P <* 10^*−*40^). The presence of central genes such as *SON, SCAF8*, and *SNRPF* suggests dysregulation of the pre-mRNA splicing machinery under APAP-induced stress.[10]
- **Extracellular Matrix and Tissue Architecture (Cluster 36)**: This cluster (*n* = 9) demonstrated enrichment in extracellular matrix organization (GO:0030198, *P <* 10^*−*20^), involving genes such as *MMP2, COL1A1*, and *COL1A2*, indicating structural remodeling of the liver microenvironment.[11, 12]
- **Hepatic Synthesis and Plasma Transport (Cluster 59)**: This module (*n* = 5) includes apolipoproteins (*APOA1, APOE, APOC3*) and *HP*, associated with lipid transport and systemic homeostasis, reflecting impaired liver function.[13]

**Figure 5.**
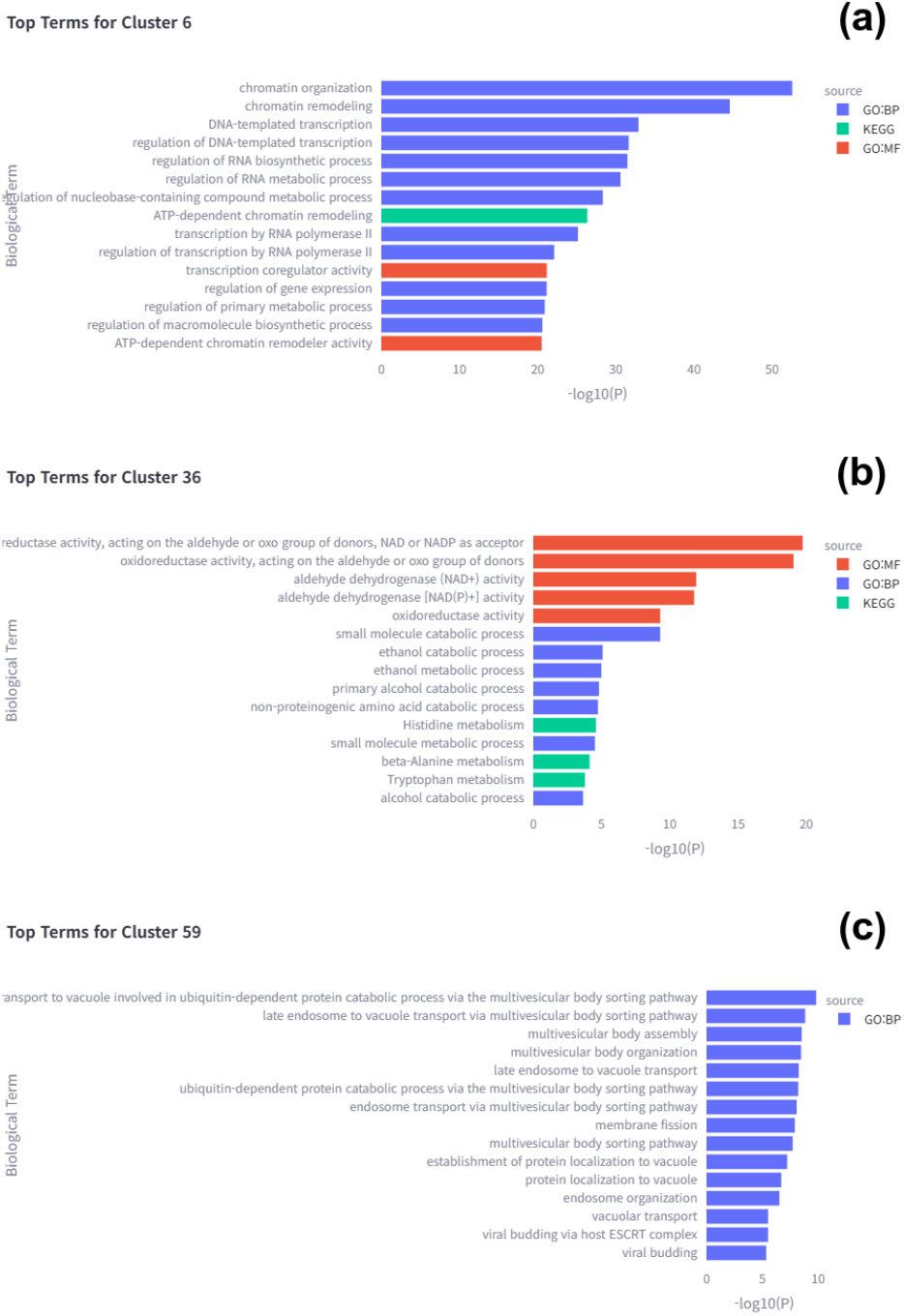
Functional enrichment of representative clusters identified by the hierarchy-aware model. (A) Cluster 6 (spliceosome), (B) Cluster 36 (extracellular matrix organization), and (C) Cluster 59 (lipid transport and synthesis).

### 3.6 Detection of Regulatory Hubs

In addition to large functional modules, the hierarchy-aware approach identified small, highly specific regulatory clusters. Notably, clusters 97 and 99, each containing three genes, revealed key regulatory hubs.

Cluster 97 (Histone modification, *P <* 10^*−*20^) includes methyltransferases such as *KMT2A* and *ASH1L*, while Cluster 99 (EP300/CREBBP complex, *P <* 10^*−*12^) highlights transcriptional co-activators involved in gene regulation.

The identification of these small but highly significant clusters demonstrates the sensitivity of the proposed method in detecting regulatory modules that may be overlooked in conventional clustering approaches.

## 4 Discussion

### 4.1 Impact of Hierarchy-aware Kernel on Functional Coherence

Our results demonstrate a substantial improvement in functional coherence, with the proposed model achieving a 33.8-fold increase compared to the expression-only baseline. While the baseline clustering produced weak enrichment signals, the hierarchy-aware model consistently identified highly significant biological modules. This improvement indicates that incorporating biological lineage through the HGNC hierarchy effectively reduces noise in transcriptomic data and aligns gene clusters with established biological pathways.

Importantly, this performance gain was achieved without modifying the downstream clustering algorithm. By maintaining an identical clustering framework across both configurations, we isolated the contribution of the hierarchy-aware similarity definition. This confirms that the improvement arises from redefining gene–gene relationships using biological priors, rather than from algorithmic tuning.

While multi-step propagation could potentially capture higher-order relationships, it may also introduce oversmoothing, leading to loss of discriminative structure.[14, 15] The single-step hyperdiffusion scheme used in this study therefore represents a balance between capturing hierarchical proximity and preserving functional specificity.

### 4.2 Mechanistic Insights from Hierarchy-aware Clustering

A central challenge in toxicogenomics is translating large lists of differentially expressed genes into interpretable biological mechanisms[16]. Conventional clustering approaches based solely on expression similarity [17] may group genes with similar magnitudes but unrelated functions.

In contrast, the proposed approach enforces functional coherence through the HGNC hierarchy. As demonstrated in the Results, clusters such as RNA processing (Cluster 6) and extracellular matrix organization (Cluster 36) correspond to well-defined biological processes. This structured grouping enables clearer interpretation of the cellular response to APAP-induced stress and facilitates reconstruction of the underlying mode of action.

### 4.3 Multi-scale Organization of Toxicity Responses

The identified clusters collectively reveal a hierarchical progression of hepatotoxicity. Early disruption of nuclear processes, reflected by spliceosome enrichment, is followed by structural remodeling of the extracellular matrix and eventual impairment of systemic functions such as lipid transport.

This progression highlights the ability of the hierarchy-aware framework to capture biological organization across multiple scales, from intracellular regulation to tissue-level and systemic responses. Such multi-level representation is difficult to achieve using expression-only approaches.

### 4.4 Interpretability through Gene-family Context

The proposed platform extends traditional list-based analysis by incorporating gene-family context into both clustering and search functionalities. Instead of presenting genes as isolated entities, the framework situates them within their hierarchical relationships.

The ALDH family example illustrates this advantage. Despite large differences in expression magnitude, genes within the same functional lineage were grouped together, preserving their shared biological role. This demonstrates that lineage-aware similarity can overcome variability in expression and improve interpretability.

Furthermore, the integration of same-cluster and hierarchy-based neighboring genes provides complementary perspectives on functional relationships, enabling users to explore both local cluster structure and broader biological context.

### 4.5 Detection of Fine-grained Regulatory Modules

In addition to large functional clusters, the method identifies small but highly significant regulatory modules. Clusters containing only a few genes, such as those associated with histone modification or transcriptional co-activation, were consistently detected with strong statistical support.

These results suggest that the hierarchy-aware approach is sensitive to tightly connected regulatory units that may be overlooked in conventional analyses. Such fine-grained modules are particularly relevant for understanding transcriptional regulation and epigenetic responses under stress conditions.

### 4.6 Limitations and Future Directions

The current framework depends on the completeness of HGNC gene family annotations. Genes without assigned families or with complex multi-family memberships may require further refinement. Future work will focus on integrating additional biological networks, such as protein–protein interactions, and extending the framework to cross-species analyses.

